# Inhibitory Circuit Compensations in Female and Male Mice: Increased Synaptic Output Offsets Reduced Parvalbumin Interneuron Density

**DOI:** 10.1101/2025.05.20.655207

**Authors:** Nadia Khoshbaf Khiabanian, Dylan J Terstege, Shi Chen Xu, Bo Young Ahn, Heewon Seo, Derya Sargin, Jonathan R Epp

## Abstract

Parvalbumin inhibitory interneurons (PV-INs) are critical regulators of excitatory/inhibitory balance in the cortex, and their dysfunction has been observed in various neurological disorders. Despite increasing recognition of sex differences in brain function, little is known about how PV-INs differ between males and females under healthy conditions. Previous work has pointed to sex differences in PV-IN vulnerability in disease and injury models. Here, we investigated sex differences in PV-IN characteristics, connectivity, and function in the retrosplenial cortex (RSC) of healthy mice. We found that female mice have significantly fewer PV-INs in the RSC compared to males, yet exhibit comparable memory induced neuronal activation (fos expression). Despite their lower numbers, female PV-INs displayed greater synaptic connectivity, as evidenced by increased synaptotagmin-2 (Syt-2) puncta per PV-IN and higher axonal bouton density. Additionally, fewer female PV-INs were surrounded by perineuronal nets (PNNs), suggesting greater plasticity in female inhibitory networks. From *ex vivo* slice electrophysiology recordings we observed greater excitability in female PV-INs compared to male PV-INs and, a reduced incidence of IPSCs. These findings indicate that female mice may compensate for reduced PV-IN numbers through enhanced synaptic output, preserving inhibitory function in the RSC. Finally, using spatial transcriptomic profiling of PV-INs we observed a number of differentially expressed genes that are consistent with the observed structural and functional differences between female and male PV-INs. Understanding these sex-specific inhibitory mechanisms is crucial for developing more targeted interventions for conditions involving PV-IN impairment and for understanding sex specific vulnerabilities to certain conditions such as Alzheimer’s disease.

## Introduction

Balance between excitation and inhibition within neural circuits is fundamental for cognitive functions such as learning and memory^1–3^. While much research has focused on excitatory neurons, inhibitory interneurons play an equally vital role in shaping neural activity and maintaining network stability^4–6^. Parvalbumin-expressing inhibitory interneurons (PV-INs) are particularly crucial for regulating high-frequency network oscillations and preserving excitatory/inhibitory (E/I) balance^7–9^. Disruptions in this balance are implicated in several neurological and psychiatric disorders, including epilepsy, schizophrenia, and Alzheimer’s disease (AD)^10–12^.

PV-INs are a subclass of fast-spiking GABAergic neurons that exert strong inhibitory control over excitatory neurons. These interneurons are critical for synchronizing network oscillations, particularly in the gamma frequency range (∼30–100 Hz), which is associated with cognitive processes including sensory integration, attention, and working memory^13–15^. Their high metabolic demands and extensive synaptic connectivity makes PV-INs vulnerable to insults, including oxidative stress and neurodegenerative processes^16^. Recently, the retrosplenial cortex (RSC) has emerged as an important region for investigating PV-IN function due to its high PV-IN density and role in spatial and episodic memory^17–21^.

Sex differences in PV-IN characteristics may contribute to differential susceptibility to neurological disorders^22,23^. For example, in mouse models of AD, RSC PV-INs show extensive deficits in structure and function in female mice, months earlier than male mice^21^. Research on PV-IN function has historically relied primarily on male subjects, leading to gaps in our understanding of potential sex-specific differences in inhibitory network dynamics. Given that females exhibit different vulnerabilities in disorders such as AD and schizophrenia, investigating baseline sex differences in PV-IN properties is essential for understanding their implications in disease susceptibility and progression^24,25^.

Here, we address this knowledge gap by examining sex differences in PV-IN functional and structural characteristics within the RSC of healthy mice. Our results indicate that female and male mice display differences in the organization of the cortical inhibitory network. These differences may contribute to differential resiliency and/or susceptibility in neurodegenerative contexts.

## Results

### Female mice have significantly fewer PV-INs in the RSC compared to male mice

PV-INs in the RSC are critical for cognitive function and are particularly vulnerable to AD pathology in female mice^21^. Despite the identification of sex differences in these cells under pathological conditions, there is currently a limited number of studies that have explicitly examined the potential for sex differences in RSC PV-INs under physiological conditions. Therefore, we first examined whether sex differences exist in the density of PV-INs within the RSC of 5-month-old male and female mice. Our analysis revealed a significantly lower density of PV-INs in female mice compared to males (Fig. 1A,B). To determine whether this sex difference was specific to the RSC, we extended our analysis to eight additional cortical regions. Across all regions examined, females consistently had significantly fewer PV-INs than males, with no significant interaction between sex and cortical region (Fig. S1A). These results suggest that the observed sex difference in PV-IN density in the RSC can be applied more broadly across cortical regions.

**Figure 1.**
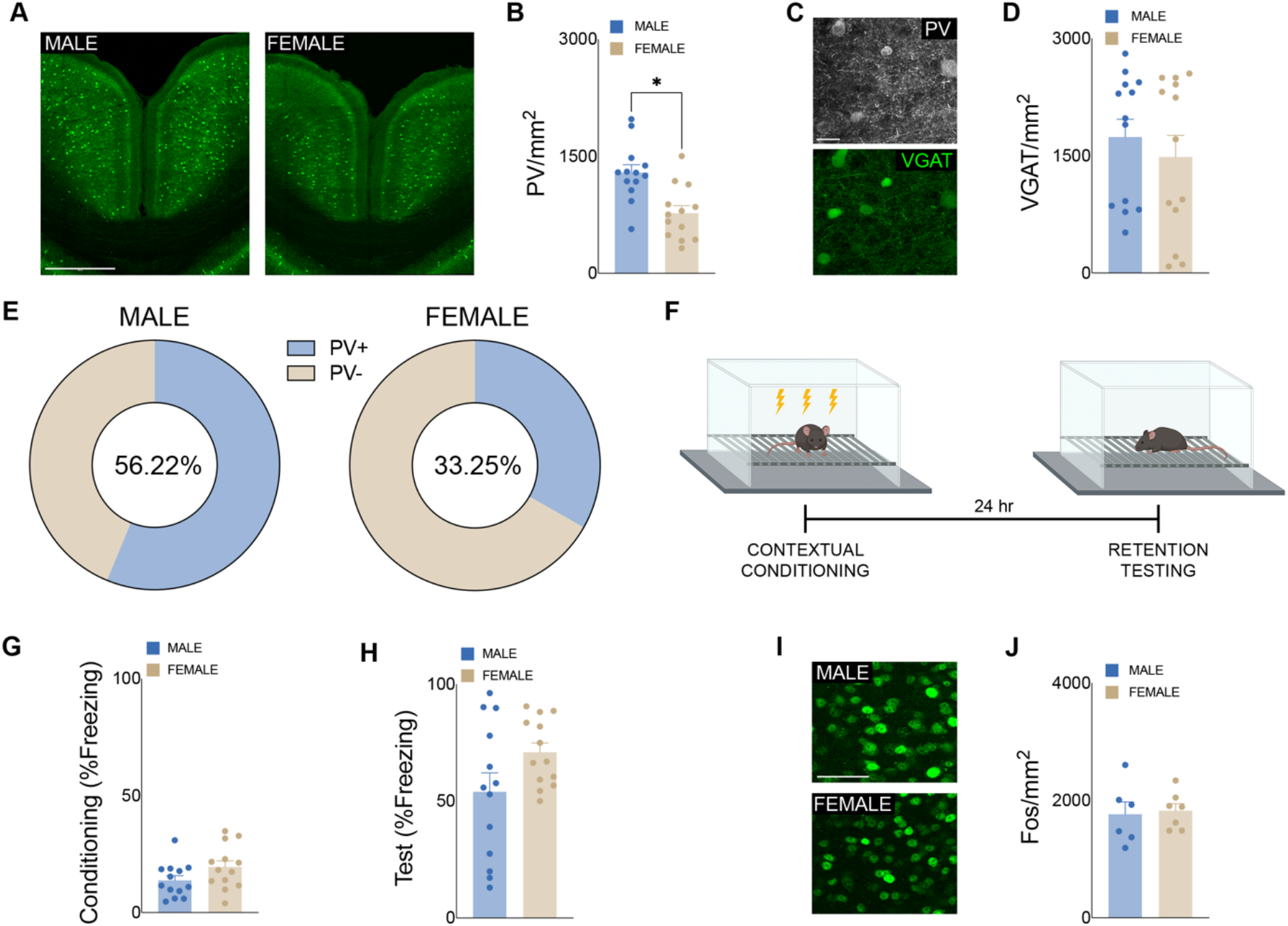
Decreased density of PV-INs in the RSC of female mice. (A) PV in the RSC of male and female mice. Scale bar represents 500 μm. (B) Decreased PV-IN density across the RSC of female (*n* = 13) compared to male mice (*n* = 13). *P* = 0.0011, two-tailed t-test. Cohen’s d = -1.46. (C) PV and VGAT labelling in the RSC. Scale bar represents 20 μm. (D) Density of VGAT across the RSC of female (*n* = 13) and male (*n* = 13) mice. Cohen’s d = -0.279. (E) Percentage of VGAT+ cells in the RSC which were PV+ in male and female mice. (F) Contextual fear conditioning protocol. (G – H) No differences in freezing behavior were observed between male (*n* = 13) and female (*n* = 13) mice across (G) conditioning or (H) test phases of contextual fear conditioning. Male conditioning vs. test Cohen’s d = 1.89. Female conditioning vs. test Cohen’s d = 4.23. (I) Fos staining in the RSC of male and female mice. Scale bar represents 50 μm. (J) Fos density across the RSC of female (*n* = 7) and male (*n* = 6) mice. Cohen’s d = 0.802.

### Total inhibitory neuron numbers are equivalent between sexes

To determine whether the reduction in PV-INs reflects a global decrease in inhibitory neurons, we quantified the overall density of inhibitory interneurons expressing VGAT (VGAT+ cells), a pan-GABAergic marker. Unlike PV-INs, no significant sex differences were observed in the density of VGAT+ cells within the RSC (Fig. 1C,D) or other cortical regions (Fig. S1B).

Further analysis revealed that PV-INs comprise a significantly smaller proportion of the total VGAT+ inhibitory interneuron population in female mice (∼33%) compared to males (∼56%; Fig. 1E). This suggests that while the absolute number of inhibitory interneurons is comparable between sexes, females have fewer PV-INs, potentially compensated for by other inhibitory subpopulations.

### Female and male mice exhibit equivalent contextual fear conditioning memory and RSC activation

To assess whether differences in PV-INs between sexes influence neuronal activation in the RSC during memory retrieval, we conducted a contextual fear conditioning (CFC) task and measured fos expression, an immediate early gene marker of neuronal activity (Fig. 1F-J). While freezing behavior was recorded as a standard indicator of memory retention, the primary goal of this experiment was to determine whether differences in PV-INs would translate into measurable differences in RSC activation during memory retrieval. During the training phase, male and female mice exhibited comparable freezing behavior, indicating similar fear acquisition (Fig. 1G). Likewise, during the 24-hour retention test, no significant sex differences were observed in freezing behavior, although a non-significant trend toward higher freezing was observed in females (Fig. 1H). Since this trend was evident in both training and retention, it is likely related to baseline locomotion or freezing thresholds rather than memory ability *per se*.

To examine neuronal activation in the RSC, we quantified fos expression 90 minutes after the retention session (Fig. 1I,J). No significant sex differences were observed in fos expression, indicating that male and female mice engaged the RSC to a similar extent during CFC retrieval. This finding suggests that differences in PV-INs between sexes do not inherently lead to differences in overall RSC activation, supporting the idea that alternative mechanisms may allow female and male mice to maintain normal inhibitory function despite their differences in PV-IN density.

### Females exhibit greater PV-IN synaptic connectivity despite fewer PV-INs

To assess sex differences in connectivity of PV-INs, we quantified the density of Synaptotagmin-2 (Syt-2)-positive presynaptic terminals in the RSC, given that Syt-2 is a marker of PV-IN synapses. Despite female mice having fewer PV-INs, the density of Syt-2 puncta across the RSC did not differ by sex (Fig. 2A,B). Normalizing this density by the density of RSC PV-IN, female mice exhibited a significantly higher density of Syt-2 puncta per PV-IN compared to males (Fig. 2C). The higher ratio of Syt-2 puncta to PV-INs in the RSC indicated that each individual PV-IN in females forms more synaptic contacts. To further investigate this, we analyzed GFP-labeled PV-IN axonal boutons in the RSC using high-resolution confocal microscopy (Fig. 2D). Female mice exhibited a significantly higher density of axonal boutons per µm of axon length compared to males (Fig. 2E). This finding suggests that female PV-INs may compensate for their lower numbers by increasing their output per cell, allowing for the maintenance of similar levels of overall inhibition despite reduced PV-IN density.

**Figure 2.**
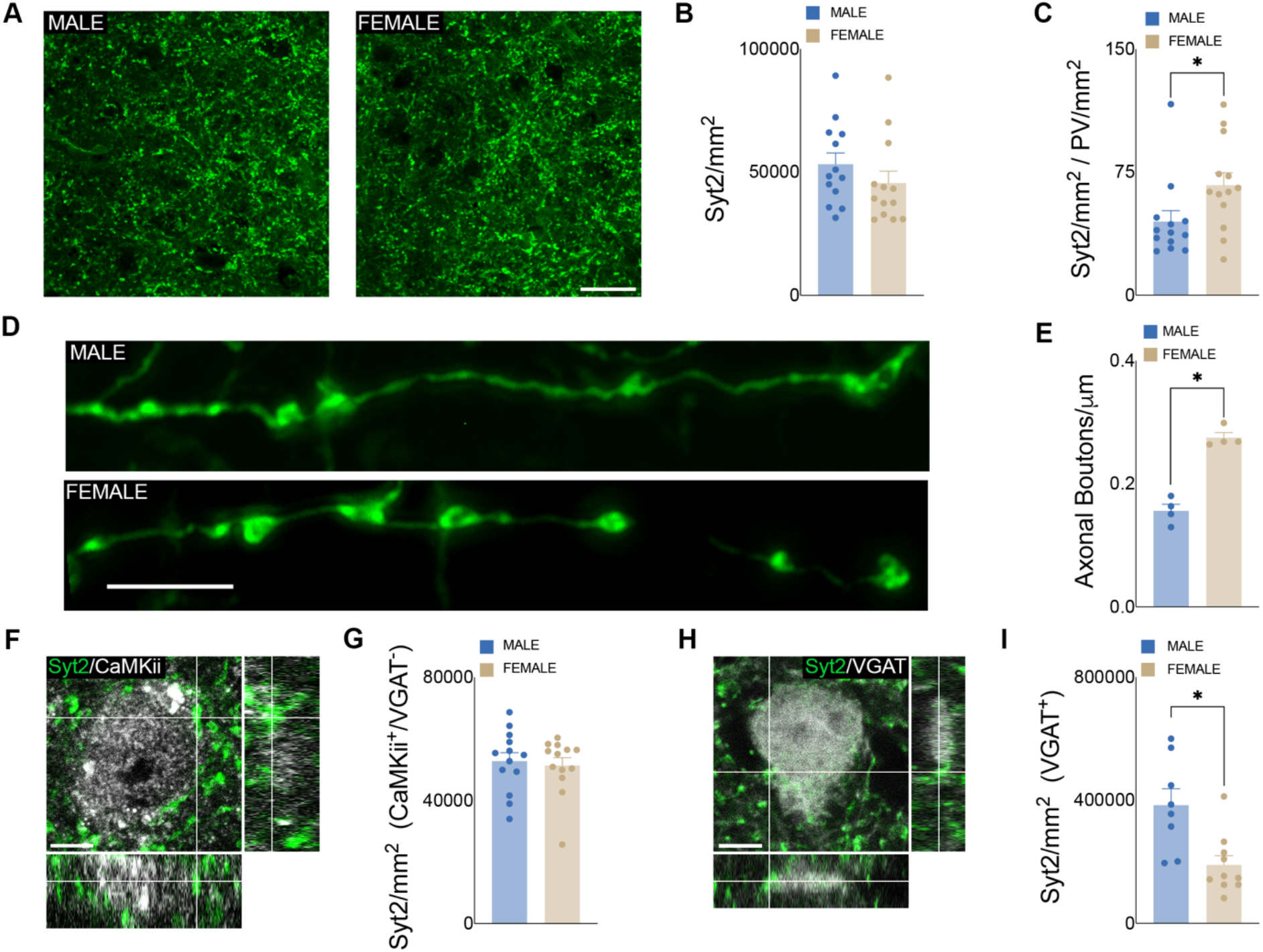
Altered synaptic connectivity profiles between RSC PV-INs of male and female mice. (A) Syt2 staining in the RSC of male and female mice. Scale bar represents 20 μm. (B) Density of syt2 labelling across the RSC of male (*n* = 13) and female (*n* = 13) mice. Cohen’s d = -0.443. (C) Normalized by the density of PV-INs, female (*n* = 13) mice had an increased density of Syt2 puncta across the RSC relative to male (*n* = 13) mice. *P* = 0.0383, two-tailed t-test. Cohen’s d = 0.86. (D) PV-IN axon labelling in the RSC of male (top) and female (bottom) mice. Scale bar represents 5 μm. (E) Female RSC PV-IN axons (*n* = 4) had a higher density of boutons than those of male (*n* = 4) mice. *P* = 0.0001, two-tailed t-test. Cohen’s d = 6.27. (F) Syt-2 (green) staining on CaMKii+ (greyscale), VGAT-cells. Scale bar represents 10 μm. (G) Syt-2 density on CaMKii+ cells. Cohen’s d = -0.146. (H) Syt-2 (green) staining on VGAT (greyscale) labelled cells. Scale bar represents 10 μm. (I) VGAT+ in the RSC of female mice (*n* = 10) were somatically innervated by fewer Syt-2 puncta than in male mice (*n* = 8). *P* = 0.0042, two-tailed t-test. Cohen’s d = -1.58.

### Female PV-INs form fewer synaptic contacts on inhibitory cell populations

While no overall differences in Syt-2 puncta density were observed across the RSC, it is important to consider the different cell types that PV-INs synapse onto. PV-INs inhibit not only excitatory cell populations, but other inhibitory neurons as well. To determine whether this increase in synaptic connectivity was specific to either inhibitory or excitatory neurons, we also quantified Syt-2 puncta density on CaMKII-expressing excitatory cells (CaMKII+/VGAT-) and VGAT-expressing inhibitory neurons. Interestingly, the density of Syt-2 puncta on excitatory neurons did not differ between sexes (Fig. 2F,G). However, the density of Syt-2 puncta was significantly lower around VGAT-labelled inhibitory populations in female mice (Fig. 2H,I). These results suggest that in female mice, the increased density of Syt-2 puncta per PV-IN are preferentially synapsing onto excitatory cell populations, with decreased inhibitory control of inhibitory populations.

### Altered expression of inhibitory synapse and dendritic markers in female and male PV-INs

To further elucidate factors driving the sex differences in RSC PV-INs, we conducted cell-type-specific spatial transcriptomic analyses using the Nanostring GeoMx Digital Spatial Profiler (DSP) platform. This process incorporates immunohistochemistry allowing for the transcriptional profiling of PV positive neurons (PV+/NeuN+; Fig. 3A) and PV negative neurons (PV-/NeuN+; Fig. S2) across the RSC of male and female mice.

**Figure 3.**
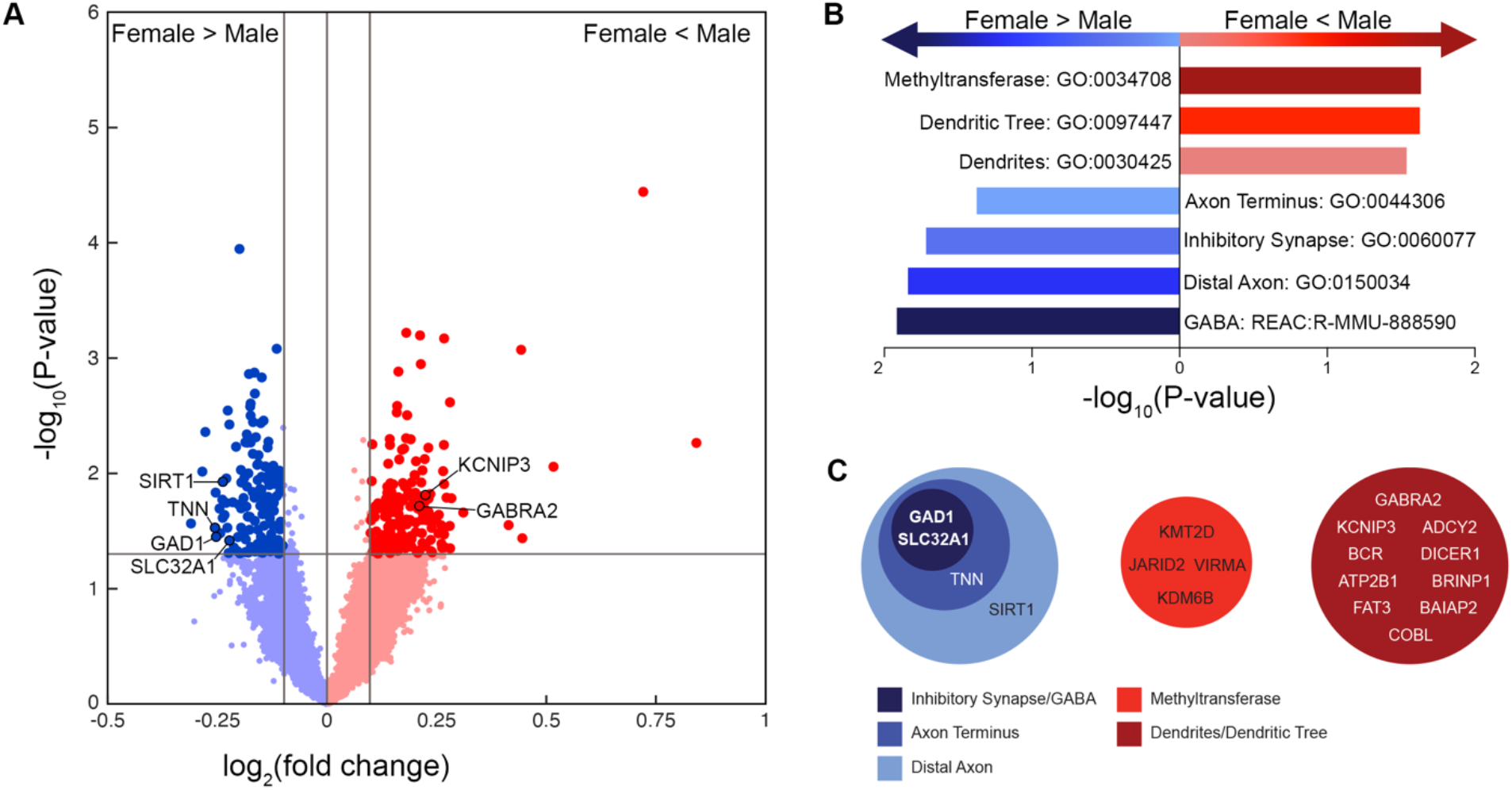
Differential gene expression profiles between RSC PV-INs of male and female mice. (A) Differentially expressed genes of RSC PV-INs. Dark blue, significantly higher expression in female mice (*n* = 3); dark red, significantly higher expression in male mice (*n* = 4); lighter colour, non-significant differential expression. (B) Top gene ontology (GO) and Reactome (REAC) terms, by fold enrichment, Blue, greater enrichment in female PV-INs than in male PV-INs; red, greater enrichment in male PV-INs than in female PV-INs. (C) Schematic highlighting differentially expressed genes involved in the differentially enriched GO and REAC terms. Overlapping circles denote differentially expressed genes which are involved in multiple processes. Blue, greater enrichment in female PV-INs than in male PV-INs; red, greater enrichment in male PV-INs than in female PV-INs.

Several themes emerged during gene ontology analyses. Male PV-INs had greater expression of genes related to methyltransferase activity, dendrites, and the outgrowth of dendritic trees, while female PV-INs showed increased expression of genes associated with axon terminus function and the formation of inhibitory, GABAergic, synapses (Fig. 3B). Looking more closely at the genes driving these changes in gene ontology analyses, the upregulation of *Gad1*and *Slc32a1* was critically involved in all functions which were increased in female mice (Fig. 3C). *Gad1* encodes the protein glutamate decarboxylase 1, an enzyme critical for the synthesis of GABA from glutamate. *Slc32a1* encodes the vesicular GABA transporter – VGAT – and is implicated in the loading of GABA into synaptic vesicles. Together, these results suggest a heightened importance of efficient GABA release in the maintenance of female-specific RSC PV-IN dynamics.

### Female PV-INs exhibit increased excitability compared to male PV-INs

*Gabra2* and *Kcnip3* which are differentially regulated in male and female PV-INs, encode proteins crucially involved in the function of the GABA-A receptor and voltage gated potassium channels, respectively. These proteins have critical roles in neuronal excitability suggesting potential sex differences in the physiological properties of PV-INs across the RSC. To investigate the functional impact of the observed differences in male and female PV-INs, we performed whole-cell patch-clamp electrophysiology on RSC slices. Current-clamp recordings revealed increased intrinsic excitability among PV-INs in the RSC of female mice, compared with PV-INs of males (Fig. 4A,B). Underlying these differences in excitability, the PV-INs in the RSC of female mice had a significantly lower spike threshold (Fig. 4C) without any differences in resting membrane potential (Fig. 4D) or other key intrinsic membrane properties (Supplemental Table S1). Coinciding with this increase in excitability, we found that the frequency of spontaneous inhibitory postsynaptic currents (sIPSCs) acting on these cells was significantly reduced in females (Fig. 4E,F). Together, these data suggest a presynaptic decrease in gross inhibitory output to PV-INs along with an increased PV-IN intrinsic excitability in the female RSC when compared to males.

**Figure 4.**
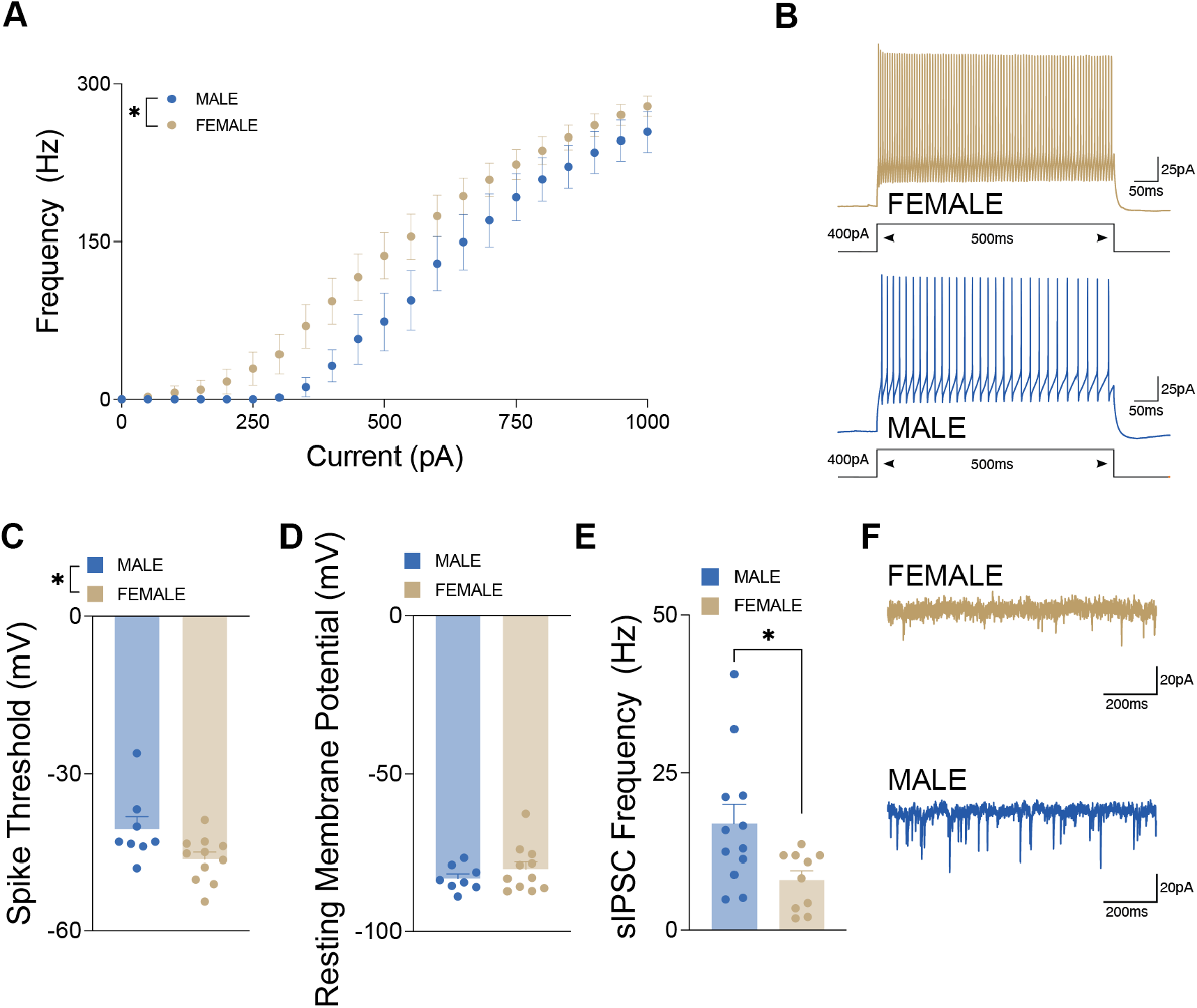
Decreased inhibitory tone across the RSC of female mice. (A) Input-output curve showing spike frequency (Hz) of pyramidal cells in male (*N* mice = 4; *n* cells = 9) and female (*N* mice = 4; *n* cells = 12) mice in response to a series of depolarization current (pA) injections. *P* < 0.0001, two-way ANOVA. (B) Representative current-clamp traces of RSC PV-INs from a female (top) and a male mouse (bottom). (C) Lower PV-IN spike threshold in female (*N* mice = 4; *n* cells = 11) mice compared to male (*N* mice = 4; *n* cells = 8) mice. *P* = 0.0358, two-tailed t-test. Cohen’s d = -1.06. (D) Resting membrane potential of RSC PV-INs in male (*N* mice = 4; *n* cells = 8) and female (*N* mice = 4; *n* cells = 11) mice. (E) Lower sIPSC frequency in female PV-INs (*N* mice = 6; *n* cells = 10) compared to male PV-INs (*N* mice = 6; *n* cells = 12). *P* = 0.0231, two-tailed t-test. Cohen’s d = -1.05. (F) Representative sIPSC traces from RSC PV-INs of a female (top) and a male (bottom) mouse.

## Discussion

This study provides compelling evidence of distinct structural and functional characteristics between male and female brains. Our findings demonstrate that while female mice have significantly fewer PV-INs compared to males, this reduction does not translate to impairments in contextual fear memory retrieval or overall RSC activity, as measured by fos expression. Instead, female PV-INs exhibit increased synaptic connectivity, axonal bouton density and excitability, suggesting alternative mechanisms that maintain inhibitory function despite a lower cell count.

The increased PV-IN connectivity in female mice is particularly intriguing. Despite having fewer PV-INs, females exhibited a higher number of synaptic contacts per PV-IN and greater Syt2 expression, a marker of presynaptic inhibitory terminals^26^. This suggests that female PV-INs may compensate for their lower numbers by increasing synaptic output, potentially sustaining inhibitory control over RSC circuits. These findings challenge traditional assumptions that inhibitory network strength is solely dependent on PV-IN quantity, emphasizing that functional connectivity and plasticity also contribute to network stability.

Beyond the RSC, our results indicate that this sex difference in PV-IN density extends across multiple cortical regions, suggesting a broader, brain-wide divergence in inhibitory interneuron populations. This widespread difference may have profound implications for neurological and psychiatric conditions that exhibit sex biases, including schizophrenia, autism, epilepsy, and AD. Given that PV-IN dysfunction has been linked to hyperexcitability and network instability in these disorders, understanding how inhibitory circuits are differentially regulated in males and females could inform sex-specific therapeutic approaches.

One particularly relevant implication of these findings is in AD, where we have shown that PV-INs in the RSC are particularly vulnerable to early dysfunction^21^. Intriguingly, our data suggest that female mice might be more resilient under normal conditions due to enhanced PV-IN connectivity. However, this connectivity-dependent compensation may render female PV-INs more susceptible to age-related or pathological insults that result in the loss of PV-INs. In the female brain, loss of each PV-IN would lead to the loss of approximately twice as many synaptic contacts.

In the current study, interpretations of inhibitory neuronal populations were limited to PV-INs. The lack of differences in the density of VGAT+ cells, despite the decrease in PV-INs, suggests an increased density of other GABAergic subpopulations across the cortex of female mice. However, it is important to consider the distinct functions and capabilities of these GABAergic subpopulations. Particularly, the calcium binding efficiency of the parvalbumin protein itself, distinct patterns of synaptic connectivity, and the high basal firing rate of these cells. Therefore, it is unlikely that other GABAergic populations fully compensate for the decreased density of cortical PV-INs of female mice and that these circuit-level compensations are more likely to be reflected in the expanded network of synaptic connections formed by female PV-INs.

In conclusion, our findings underscore the need for greater inclusion of both sexes in preclinical and clinical research, particularly in studies of neurological and psychiatric disorders where sex-specific vulnerabilities and resilience mechanisms may shape disease onset and progression. We show here, differential organization of the cortical PV-IN system in males and females which, both appear to provide effective inhibition but may confer different vulnerability to disease states. Future research should further explore the molecular and physiological basis of these sex differences, particularly how they interact with aging, hormonal fluctuations, and disease states, to inform more effective, sex-tailored therapeutic strategies.

## Resource availability

### Lead contact

Further information and requests for resources and reagents should be directed to the lead contact, Jonathan Epp.

## Materials availability

This study did not generate new unique materials.

## Data and code availability

The datasets generated during this study are available upon request from the lead contact. No new code was generated for data analysis.

## Acknowledgements

Funding for this study was provided by a Canadian Foundation for Innovation (CFI) Grant (#38160) to J.R.E. Additional funding was provided by a CFI John R. Evans Leaders Fund Grant (#42065) to D.S. Both D.S. and J.R.E. received additional funding from an Alberta Children’s Hospital Research Institute Child Health and Wellness Grand Challenge Seedling Award and a Women’s Brain Health Initiative Grant in partnership with Brain Canada (#5542). D.J.T. received a doctoral fellowship from NSERC (CGS D). Transcriptomic analyses were performed with assistance from the Cumming School of Medicine Applied Spatial Omics Centre. The znp-1 monoclonal antibody (AB_2315626) developed by Bill Trevarrow and colleagues was obtained from the Developmental Studies Hybridoma Bank, created by the NICHD of the NIH and maintained by the University of Iowa, Department of Biological Sciences, Iowa City, IA, USA.

## Author contributions

N.K. contributed to conceptualization/study design, histological procedures, data analysis, figure generation, and manuscript preparation; D.J.T. contributed to histological procedures/analyses, data analysis, figure generation, manuscript preparation; S.X. performed the electrophysiological recordings, contributed to data analysis, and manuscript preparation; B.A. advised on transcriptomic experimental design and contributed to the preparation of transcriptomic experiments; H.S. advised on transcriptomic experimental design and contributed to the analysis of transcriptomic data; D.S contributed to conceptualization/study design, supervision, direction of data analysis, manuscript preparation, and funding support acquisition; J.R.E. contributed to conceptualization/study design, supervision, direction of data analysis, manuscript preparation, and funding support acquisition.

## Methods

### Experimental model and subject details

All experiments were conducted using male and female mice aged five months at the time of testing. Mice were housed in standard cages in groups of three to five per cage, with ad libitum access to food and water. The housing conditions were maintained on a 12-hour light-dark cycle with controlled temperature and humidity. All experimental procedures were performed during the light phase.

Two distinct transgenic mouse lines were used for labeling inhibitory interneurons. VGAT-tdTomato mice were generated by crossing VGAT-Cre (Slc32a1tm2 (cre) Lowl/J, Jackson Laboratory, strain #016962) with Rosa-tdTomato mice (B6.Cg-Gt[ROSA]26Sortm9[CAG-tdTomato]Hze/J, strain #007909). PV-Cre mice (B6 Pvalb-IRES-Cre, Jackson Laboratory, strain #017320) were bred to selectively label PV-expressing neurons. Experimental mice were bred at the University of Calgary’s Clara Christie Centre for Mouse Genomics.

All experimental procedures were conducted in accordance with the Canadian Council on Animal Care (CCAC) guidelines and approved by the University of Calgary Animal Care Committee.

## Method details

### Contextual fear conditioning (CFC)

To assess memory function, mice underwent contextual fear conditioning (CFC) using sound-attenuated chambers (Ugo Basile, Italy, 17 × 17 × 24.7 cm) equipped with a metal grid floor for shock delivery. Mice were habituated to handling for three consecutive days before testing. During the training session, each mouse was placed in the chamber and allowed to explore for two minutes before receiving three 0.5 mA foot shocks (each lasting two seconds), separated by one-minute intervals. Mice were removed from the chamber one minute after the final shock and returned to their home cages. Twenty-four hours later, mice were placed back into the same context for five minutes with no shock, and freezing behavior was recorded as a measure of memory retention. Freezing behavior was analyzed using an overhead infrared camera and ANY-maze tracking software (Stoelting, Wood Dale, IL, USA).

### Tissue processing and immunohistochemistry

Mice were transcardially perfused 90 minutes after the retention test with ice-cold 0.1 M phosphate-buffered saline (PBS), followed by 4% paraformaldehyde (PFA) in PBS. Brains were post-fixed in 4% PFA for 24 hours and then cryoprotected in 30% sucrose solution at 4°C for three to five days. Coronal brain sections (50 µm thick) were obtained using a Leica CM1950 cryostat (Leica Microsystems, Germany) and stored at -20°C in an antifreeze solution containing 30% ethylene glycol and 20% glycerol in 0.1 M PBS.

### Fos immunolabeling

To assess neuronal activation, fos immunolabeling was performed. Free-floating brain sections were washed in 0.1 M PBS (three 10-minute washes) before incubation in primary antibody solution containing 1:2000 anti-c-Fos (RPCA-C-FOS, Encor Biology, AB_2572236), 3% normal donkey serum, and 0.3% Triton X-100 in 0.1 M PBS. Sections were incubated for 48 hours at room temperature. After washing, sections were incubated for 24 hours at room temperature with 1:500 Alexa Fluor 647-conjugated secondary antibody (anti-rabbit, Jackson ImmunoResearch). Finally, sections were counterstained with DAPI (1:1000 in PBS, 20 minutes) before coverslipping with PVA-DABCO anti-fade medium.

### Parvalbumin (PV) immunolabeling

PV-expressing neurons were labeled using 1:5000 rabbit anti-parvalbumin antibody (Invitrogen, PA1-933) in 3% normal donkey serum and 0.3% Triton-X100 in 0.1 M PBS. Sections were incubated for 24 hours at room temperature, followed by incubation with 1:500 goat anti-rabbit Alexa Fluor 647 (Jackson ImmunoResearch) for 24 hours. Sections were counterstained with DAPI, washed, and mounted using PVA-DABCO.

### Synaptotagmin-2 (Syt-2) and CaMKII immunolabeling

To assess synaptic connectivity, sections were incubated with 1:5000 anti-PV (Invitrogen, PA1-933) and 1:200 anti-Syt-2 (ZNP-1, DSHB, AB_2315626) for 48 hours at room temperature. After washing, sections were incubated with 1:500 Alexa Fluor 647-conjugated goat anti-rabbit and 1:500 Alexa Fluor 488-conjugated alpaca anti-mouse secondary antibodies (Jackson ImmunoResearch) for 24 hours.For co-labeling with excitatory neurons, sections were incubated with anti-Syt-2 and anti-CaMKII antibodies, followed by corresponding fluorophore-conjugated secondary antibodies.

### Microscopy and image analysis

Confocal imaging was conducted using an Olympus FV3000 microscope equipped with 10X (NA 0.42), 20X (NA 0.75), and 60X oil-immersion (NA 1.42) objectives. Images were analyzed using FIJI/ImageJ. To quantify cell density, Ilastik machine-learning software was used for automated cell segmentation. Ilastik was trained on a subset of manually labeled images and validated against manual counts, with an error rate of <5%. For synaptic connectivity analysis, Syt-2 puncta were identified using Ilastik segmentation and quantified within PV+ and CaMKII+ cell boundaries using FIJI.

### *Ex vivo* slice electrophysiology

Coronal slices of the RSC at 400µm thickness were obtained in ice-cold sucrose-ACSF cutting solution (254mM sucrose, 10mM D-glucose, 26Mm NaHCO_3_, 2mM CaCl_2_·2H_2_O, 2mM MgSO_4_, 3mM KCl, 1.25mM NaH_2_PO_4_) saturated with 5% CO_2_, 95% O_2_. Recovered slices were incubated for 1 hour and recorded in ACSF (128mM NaCl, 10mM D-glucose, 26mM NaHCO_3,_ 2mM CaCl_2_·2H_2_O, 2mM MgSO_4_, 3mM KCl, 1.25mM NaH_2_PO_4_) saturated with 5% CO_2_, 95% O_2_ at 30^0^C. Current-clamp recordings were performed with electrodes (3MΩ-6MΩ) containing K-gluconate internal patch solution (120mM K-gluconate, 10mM HEPES, 5mM KCl, 2mM MgCl_2_, 2mM K2-ATP, 0.4mM Na_2_-GTP, 10 mM Na2-phosphocreatine, adjusted to 7.3 with KOH). Voltage-clamp recordings of sIPSCs were obtained with high Cl^-^ patch solution (50mM K-gluconate, 10mM HEPES, 75mM KCl, 2mM MgCl_2_, 2mM K2-ATP, 0.4mM Na_2_-GTP, 10 mM Na2-phosphocreatine, adjusted to 7.3 with KOH). Cells were visualised with an Olympus BX51WI equipped with 5X (NA 0.15) and X40 water-immersion (NA 0.8) objectives with white light from an Olympus TH4-100 passed through an IR-filter and Olympus differential interference contrast mirror WI-DPMC. PV+ neurons were visualized based on tdTomato expression using CoolLED pE-300^ultra^ LED light source. Whole-cell recordings were performed using the Multiclamp 700B amplifier (Molecular Devices). Data were acquired at 20 kHz and 100 kHz (current-clamp) and low-pass filtered at 2-4 kHz. All data were acquired using pClamp11.1 Software Suite with Humsilencer and Axon Digidata1550B software. Voltage-clamp sIPSCs recordings were analyzed in SimplyFire^27^. Current-clamp input-output recordings were analyzed using Easy Electrophysiology. All other analyses were performed using pClamp11.1 software.

### GeoMX spatial transcriptomics

Following perfusion and post-fixation, brains were embedded in paraffin and sectioned at a thickness of 5 μm and mounted onto slides. Tissue was prepared for GeoMx Digital Spatial Profiler (DSP; NanoString, Inc., v2.5) as per the manufacturer’s instructions (NanoString, Inc., MAN-10150)^28^. Prior to hybridization, slides underwent antigen retrieval and RNA exposure steps using citrate buffer and 0.1 μg/mL of proteinase K treatment, respectively, to optimize accessibility of both cell markers and RNA for downstream probe hybridization. *In situ* hybridizations were performed using the GeoMx Mouse Whole Transcriptome Atlas (WTA), which targets 20,175 genes and includes 210 negative control designed against sequences absent from the mouse genome. For subsequent antibody staining, slides were incubated for 1 h in a solution of 1:100 Alexa Fluor 488 conjugated anti-NeuN (Sigma Aldrich; ABN78A4), 1:100 Alexa Fluor 594 (Abcam; ab269822) conjugated in-house to an anti-PV antibody (Invitrogen; PA1-933) and 1:50 SYTO 83 (Thermo Fisher; S11364). Following a series of washes in sodium citrate buffer solution, slides were immediately scanned at 20X using the GeoMx DSP platform.

Regions of interest (ROIs) were selected on each section (1-2 per section) to encompass the RSC. Within each ROI, fluorescently labelled PV+ and NeuN+ cells were segmented from background to define areas of illumination (AOIs) corresponding to PV-INs (PV+/NeuN+) and PV negative neurons (PV-/NeuN+). The GeoMx DSP platform then photocleaved a UV-cleavable barcode linker bound to the RNA *in situ* probes, releasing bound oligonucleotides from all nuclei within the PV-IN ROI and deposited these in the DSP collection plate. Afterwards, oligonucleotides were cleaved from the PV-/NeuN+ ROI and deposited in a separate well of the DSP collection plate. These procedures were then repeated for both cell types across all ROIs.

To uniquely index each AOI for sequencing, Illumina i5 and i7 dual-indexing primers were added to the oligonucleotide tags during PCR amplification. Libraries were purified using AMPure XP beads (Beckman Coulter), and quality was assessed using an Agilent Bioanalyzer. Concentration was determined using a Qubit fluorometer (Thermo Fisher Scientific). Sequencing was performed on an Illumina NextSeq 2000 (Illumina Inc.), targeting a sequencing depth of 100 counts/μm^2^.

Using the NanoString GeoMx NGS Pipeline (NanoString, Inc., MAN-10153), FASTQ files were processed into gene count data for each AOI. Quality control of these data was implemented on a label-by-label basis by assessing the quality of the isolated segments. Assessments incorporated the percentage of trimmed reads (>80%), the percentage of stitched reads (>80%), the percentage of aligned reads (>75%), the sequencing saturation (>50%), the area of the AOI (>10000μm^2^), and a minimum number of nuclei (>20), as per manufacture recommendations. Low performance probes were identified by dividing the geometric mean of a single probe count across all samples against the geometric mean of all the probe counts for that gene. Cases of non-specific binding were identified by defining the limit of quantification as 2 standard deviations above the geometric mean of the negative probes for each sample. If fewer than 5% of genes were detected above the limit of quantification, AOIs were removed from analyses. Following these measures, 8,113 genetic targets remained.

Raw counts were normalized using variance stabilizing transformation (DESeq2)^29^. Differential expression within PV+ and NeuN+ cells was calculated between males and females for each gene using a linear mixed effect model with the GeoMx Tools R package (v3.4.0). Changes in gene expression with *p* values <0.05 and an absolute log2 fold-change of >1.0 were defined as being differentially expressed. Volcano plots were generated using the MATLAB ‘volcanoplot’ function from the RokaiXplorer toolbox ^30^. Gene ontology (GO) and Reactome (REAC) analyses were performed using g:Profiler ^31–33^. FDR-based *p* value correction was used for all analyses.

### Quantification and statistical analysis

All statistical analyses were performed in GraphPad Prism (Version 9.4.0). Independent t-tests were used for comparisons between male and female mice. For multiple-region analyses, two-way ANOVAs were conducted with sex and brain region as factors. Hypothesis testing was supplemented with estimation statistics to report effect sizes (Cohen’s d) and 95% confidence intervals. All statistical tests were two-tailed, with significance set at p < 0.05. Data are presented as mean ± standard error of the mean (SEM).

## References

1. English, D.F., Peyrache, A., Stark, E., Roux, L., Vallentin, D., Long, M.A., and Buzsáki, G. (2014). Excitation and inhibition compete to control spiking during hippocampal ripples: intracellular study in behaving mice. J. Neurosci. 34, 16509–16517.

2. Sengupta, B., Laughlin, S.B., and Niven, J.E. (2013). Balanced excitatory and inhibitory synaptic currents promote efficient coding and metabolic efficiency. PLoS Comput. Biol. 9, e1003263.

3. Agnes, E.J., and Vogels, T.P. (2024). Co-dependent excitatory and inhibitory plasticity accounts for quick, stable and long-lasting memories in biological networks. Nat. Neurosci. 27, 964–974.

4. Ben-Ari, Y. (2014). The GABA excitatory/inhibitory developmental sequence: a personal journey. Neuroscience 279, 187–219.

5. Bocchio, M., Vorobyev, A., Sadeh, S., Brustlein, S., Dard, R., Reichinnek, S., Emiliani, V., Baude, A., Clopath, C., and Cossart, R. (2024). Functional networks of inhibitory neurons orchestrate synchrony in the hippocampus. PLoS Biol. 22, e3002837.

6. Mueller-Buehl, C., Wegrzyn, D., Bauch, J., and Faissner, A. (2023). Regulation of the E/I-balance by the neural matrisome. Front. Mol. Neurosci. 16, 1102334.

7. Ferguson, B.R., and Gao, W.-J. (2018). PV interneurons: Critical regulators of E/I balance for prefrontal cortex-dependent behavior and psychiatric disorders. Front. Neural Circuits 12. 10.3389/fncir.2018.00037.

8. Gonzalez-Burgos, G., Cho, R.Y., and Lewis, D.A. (2015). Alterations in cortical network oscillations and parvalbumin neurons in schizophrenia. Biol. Psychiatry 77, 1031–1040.

9. Canty, A.J., Dietze, J., Harvey, M., Enomoto, H., Milbrandt, J., and Ibáñez, C.F. (2009). Regionalized loss of parvalbumin interneurons in the cerebral cortex of mice with deficits in GFRalpha1 signaling. J. Neurosci. 29, 10695–10705.

10. Duma, G.M., Cuozzo, S., Wilson, L., Danieli, A., Bonanni, P., and Pellegrino, G. (2024). Excitation/Inhibition balance relates to cognitive function and gene expression in temporal lobe epilepsy: a high density EEG assessment with aperiodic exponent. Brain Commun. 6, fcae231.

11. Lányi, O., Koleszár, B., Schulze Wenning, A., Balogh, D., Engh, M.A., Horváth, A.A., Fehérvari, P., Hegyi, P., Molnár, Z., Unoka, Z., et al. (2024). Excitation/inhibition imbalance in schizophrenia: a meta-analysis of inhibitory and excitatory TMS-EMG paradigms. Schizophrenia (Heidelb.) 10, 56.

12. van Nifterick, A.M., Mulder, D., Duineveld, D.J., Diachenko, M., Scheltens, P., Stam, C.J., van Kesteren, R.E., Linkenkaer-Hansen, K., Hillebrand, A., and Gouw, A.A. (2023). Resting-state oscillations reveal disturbed excitation-inhibition ratio in Alzheimer’s disease patients. Sci. Rep. 13, 7419.

13. Hadler, M.D., Tzilivaki, A., Schmitz, D., Alle, H., and Geiger, J.R.P. (2024). Gamma oscillation plasticity is mediated via parvalbumin interneurons. Sci. Adv. 10, eadj7427.

14. Kriener, B., Hu, H., and Vervaeke, K. (2022). Parvalbumin interneuron dendrites enhance gamma oscillations. Cell Rep. 39, 110948.

15. Volman, V., Behrens, M.M., and Sejnowski, T.J. (2011). Downregulation of parvalbumin at cortical GABA synapses reduces network gamma oscillatory activity. J. Neurosci. 31, 18137–18148.

16. Ruden, J.B., Dugan, L.L., and Konradi, C. (2021). Parvalbumin interneuron vulnerability and brain disorders. Neuropsychopharmacology 46, 279–287.

17. Klimczak, P., Rizzo, A., Castillo-Gómez, E., Perez-Rando, M., Gramuntell, Y., Beltran, M., and Nacher, J. (2021). Parvalbumin interneurons and perineuronal nets in the hippocampus and retrosplenial cortex of adult male mice after early social isolation stress and perinatal NMDA receptor antagonist treatment. Front. Synaptic Neurosci. 13, 733989.

18. Park, K., Kohl, M.M., and Kwag, J. (2024). Memory encoding and retrieval by retrosplenial parvalbumin interneurons are impaired in Alzheimer’s disease model mice. Curr. Biol. 34, 434-443.e4.

19. Terstege, D.J., and Epp, J.R. (2025). Cognitive enrichment preserves retrosplenial parvalbumin density and cognitive function in female 5xFAD mice. bioRxiv, 2025.01.15.633249. 10.1101/2025.01.15.633249.

20. Hu, Y., Feng, Y., Luo, H., Zhu, X.-N., Chen, S., Yang, K., Deng, Z., Luo, M., Du, W., Wang, Q., et al. (2025). Dissociation-related behaviors in mice emerge from the inhibition of retrosplenial cortex parvalbumin interneurons. Cell Rep. 44, 115086.

21. Terstege, D.J., Ren, Y., Ahn, B.Y., Seo, H., Adigun, K., Alzheimer’s Disease Neuroimaging Initiative, Galea, L.A.M., Sargin, D., and Epp, J.R. (2025). Impaired parvalbumin interneurons in the retrosplenial cortex as the cause of sex-dependent vulnerability in Alzheimer’s disease. Sci. Adv. 11. 10.1126/sciadv.adt8976.

22. Woodward, E.M., and Coutellier, L. (2021). Age- and sex-specific effects of stress on parvalbumin interneurons in preclinical models: Relevance to sex differences in clinical neuropsychiatric and neurodevelopmental disorders. Neurosci. Biobehav. Rev. 131, 1228–1242.

23. Terstege, D.J., and Epp, J.R. (2023). Parvalbumin as a sex-specific target in Alzheimer’s disease research - A mini-review. Neurosci. Biobehav. Rev. 153, 105370.

24. Ferretti, M.T., Iulita, M.F., Cavedo, E., Chiesa, P.A., Schumacher Dimech, A., Santuccione Chadha, A., Baracchi, F., Girouard, H., Misoch, S., Giacobini, E., et al. (2018). Sex differences in Alzheimer disease - the gateway to precision medicine. Nat. Rev. Neurol. 14, 457–469.

25. Sommer, I.E., Tiihonen, J., van Mourik, A., Tanskanen, A., and Taipale, H. (2020). The clinical course of schizophrenia in women and men-a nation-wide cohort study. NPJ Schizophr. 6, 12.

26. Sommeijer, J.-P., and Levelt, C.N. (2012). Synaptotagmin-2 is a reliable marker for parvalbumin positive inhibitory boutons in the mouse visual cortex. PLoS One 7, e35323.

27. Mori, M., Rosko, A., Farnsworth, J., Carrasco, G., Broomandkhoshbacht, P., Pareja-Navarro, K., and Pejmun Haghighi, A. (2024). SimplyFire: An open-source, customizable software application for the analysis of synaptic events. eNeuro 11. 10.1523/ENEURO.0326-23.2023.

28. Merritt, C.R., Ong, G.T., Church, S.E., Barker, K., Danaher, P., Geiss, G., Hoang, M., Jung, J., Liang, Y., McKay-Fleisch, J., et al. (2020). Multiplex digital spatial profiling of proteins and RNA in fixed tissue. Nat. Biotechnol. 38, 586–599.

29. Love, M.I., Huber, W., and Anders, S. (2014). Moderated estimation of fold change and dispersion for RNA-seq data with DESeq2. Genome Biol. 15. 10.1186/s13059-014-0550-8.

30. Yılmaz, S., Tavares Pereira Lopes, F.B., Schlatzer, D., Ayati, M., Chance, M.R., and Koyutürk, M. (2024). Making Proteomics Accessible: RokaiXplorer for interactive analysis of phospho-proteomic data. Bioinform. Adv., vbae077.

31. Ashburner, M., Ball, C.A., Blake, J.A., Botstein, D., Butler, H., Cherry, J.M., Davis, A.P., Dolinski, K., Dwight, S.S., Eppig, J.T., et al. (2000). Gene ontology: tool for the unification of biology. The Gene Ontology Consortium. Nat. Genet. 25, 25–29.

32. Milacic, M., Beavers, D., Conley, P., Gong, C., Gillespie, M., Griss, J., Haw, R., Jassal, B., Matthews, L., May, B., et al. (2024). The reactome pathway knowledgebase 2024. Nucleic Acids Res. 52, D672–D678.

33. Kolberg, L., Raudvere, U., Kuzmin, I., Adler, P., Vilo, J., and Peterson, H. (2023). g:Profiler-interoperable web service for functional enrichment analysis and gene identifier mapping (2023 update). Nucleic Acids Res. 51, W207–W212.

