## Supplementary material for "Inhibitory Circuit Compensations in Female and Male Mice: Increased Synaptic Output Offsets Reduced Parvalbumin Interneuron Density": Document S1

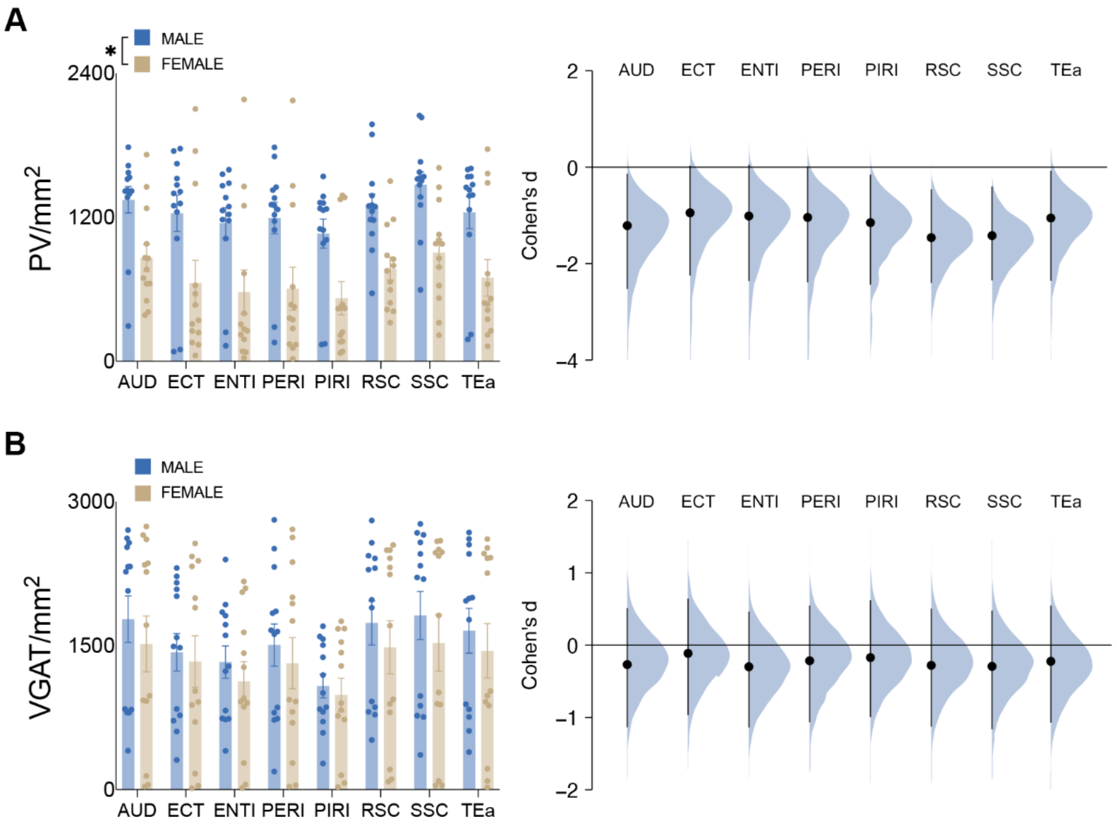


**Figure S1. Sex differences in PV-IN density across the cortex. Related to Figure 1.**

(A) Density of PV-INs across the auditory cortex (AUD), ectorhinal cortex (ECT), lateral entorhinal cortex (ENTl), perirhinal cortex (PERI), piriform cortex (PIRI), RSC, somatosensory cortex (SSC), and temporal association area (TEa). From this cortical examination, male mice ($n$ = 13) had a higher density of PV-INs than female mice ($n$ = 13). $P < 0.0001$, two-way repeated-measures ANOVA. Cohen's d: AUD = -1.21, ECT = -0.944, ENTl = -1.01, PERI = -1.04, PIRI = -1.15, RSC = -1.46, SSC = -1.41, TEa = -1.05.

(B) Density of VGAT-labelled cells across the AUD, ECT, ENTl, PERI, PIRI, RSC, SSC, and TEa. Cohen's d: AUD = -0.267, ECT = -0.114, ENTl = -0.299, PERI = -0.215, PIRI = -0.170, RSC = -0.279, SSC = -0.293, TEa = -0.224.

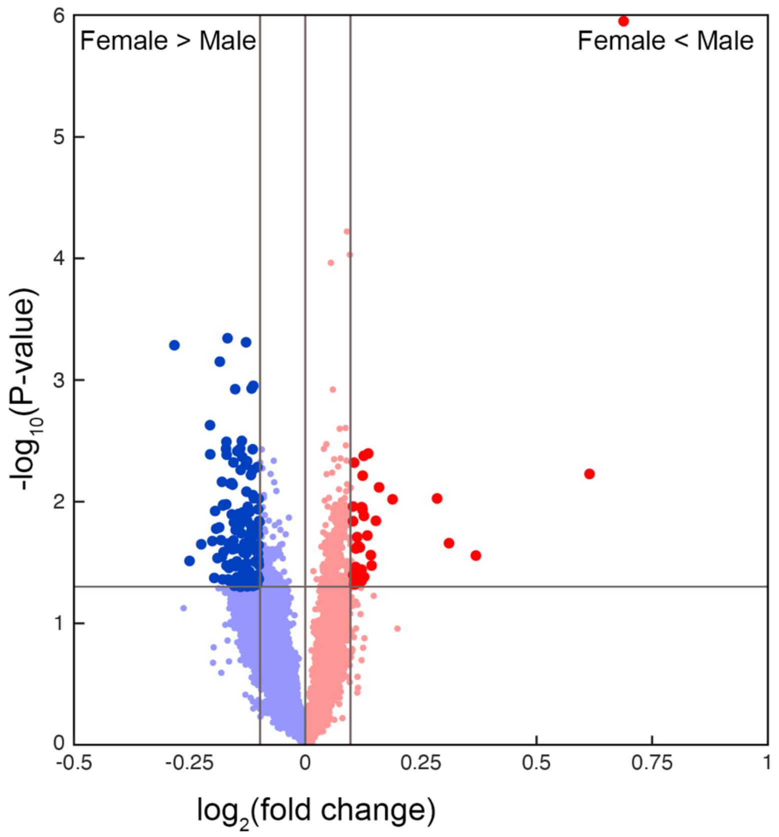


**Figure S2. Differential gene expression in non-PV-IN neuronal populations. Related to Figure 3.**

Volcano plot of differentially expressed genes of RSC neurons which did not express PV (PV-/NeuN+). Dark blue, significantly higher expression in female mice (*n* = 3); dark red, significantly higher expression in male mice (*n* = 4); lighter colour, non-significant differential expression.

**Table S1. Membrane properties of RSC PV-INs in male and female mice. Related to Figure 4.** Membrane properties were assessed from 8 RSC PV-INs from male mice and 11 RSC PV-INs from female mice. All statistical comparisons presented as unpaired Student's *t* tests.

| Property | Males Mean ± SEM | Females Mean ± SEM | *P* value | *T* stat | Cohen's D |
|---|---|---|---|---|---|
| Capacitance (pF) | 60.56 ± 5.130 | 55.61 ± 4.694 | P=0.4911 | $t_{17}$=0.7037 | -0.327 |
| Membrane Tau Constant (µs) | 464.6 ± 34.38 | 451.5 ± 17.06 | P=0.7157 | $t_{17}$=0.3704 | -0.172 |
| Input Resistance (MΩ) | 77.96 ± 3.860 | 85.63 ± 5.717 | P=0.3208 | $t_{17}$=1.023 | 0.475 |
| Spike Amplitude (mV) | 62.21 ± 3.689 | 63.99 ± 2.788 | P=0.7005 | $t_{17}$=0.3913 | 0.182 |
